# Dynamic behavior of cell-free mitochondrial DNA in human saliva

**DOI:** 10.1101/2021.09.15.460321

**Authors:** Caroline Trumpff, Shannon Rausser, Rachel Haahr, Kalpita R. Karan, Gilles Gouspillou, Eli Puterman, Clemens Kirschbaum, Martin Picard

## Abstract

Mitochondria contain their own genome that can be released in multiple biofluids such as blood and cerebrospinal fluid, as cell-free mitochondrial DNA (cf-mtDNA). In clinical studies, single measures of blood cf-mtDNA predict mortality, and higher cf-mtDNA levels are associated with mental and physical stress. However, the dynamics of cf-mtDNA has not been defined, and whether it can be measured non-invasively like other neuroendocrine markers in saliva has not been examined. Here we report cf-mtDNA in human saliva and establish its natural within-person dynamic behavior across multiple weeks. In a small proof-of-principle cohort of healthy adults, we first develop an approach to rapidly quantify salivary cf-mtDNA without DNA extraction, and demonstrate the existence of saliva cf-mtDNA. We then deploy this approach to perform an intensive repeated-measures analysis of two healthy men studied at 4 daily timepoints over 53-60 consecutive days (n=212-220 observations each) with parallel measures of steroid hormones, self-reported daily mood, and health-related behaviors. Salivary cf-mtDNA exhibited a robust awakening response reaching up to two orders of magnitude 30-45 minutes after awakening, varied from day-to-day, and moderately correlated with the cortisol awakening response. No consistent association with self-reported daily mood/health-related behaviors were found, although this requires further examination in more extensive studies. Dynamic variation in cf-mtDNA was inversely related with salivary interleukin 6 (IL6), inconsistent with a pro-inflammatory effect of salivary cf-mtDNA. The highly dynamic behavior of salivary cf-mtDNA opens the door to non-invasive studies examining the relevance of mtDNA signaling in relation to human health.

## A. Introduction

The maintenance of human health requires dynamic variations in key physiological systems (Sterling, 2020). One well-described example of dynamic regulation is the hypothalamic-pituitary-adrenal (HPA) axis, which leads to cortisol release. Cortisol is not only released acutely in response to physical or psychosocial stress (Caplin et al., 2021; Foley and Kirschbaum, 2010), but also exhibits a stereotypical diurnal pattern of variation where it rapidly increases during the first 30-45 min after awakening – a phenomenon termed the cortisol awakening response (CAR) – subsequently reaching its nadir at night (Stalder et al., 2016). The CAR is believed to sustain the metabolic activation associated with the morning-related rise in body temperature, increased heart rate, cognitive processes, and increased energy expenditure as the organism transitions from sleep to wakefulness (Rohleder et al., 2007; Schlotz et al., 2004). Diurnal variation also applies more broadly to sex hormones (Brambilla et al., 2009), circulating immune cell composition (Ackermann et al., 2012; Born et al., 1997; Scheiermann et al., 2012), metabolites (Ang et al., 2012; Chua et al., 2013; Dallmann et al., 2012; Kasukawa et al., 2012), and even organ size (Liu et al., 2021), illustrating the pervasiveness of dynamic variation in human physiology.

Dynamic, diurnal cortisol changes are readily detectable in saliva, an accessible biofluid widely used in clinical and epidemiological studies, including in special populations (e.g., children), where biospecimen collection is limited to non-invasive measures. Studies using salivary cortisol have demonstrated that an abnormal CAR is linked to cardiovascular, autoimmune, and psychiatric diseases (Kudielka and Kirschbaum, 2003). However, cortisol alone is not sufficient to fully explain or predict the diversity of adverse health outcomes, and one emerging contributor to psychobiological processes are the energy-producing organelle mitochondria. Mitochondria are both targets and mediators of the stress response (Picard and McEwen, 2018), and can influence broad aspects of physiology (Picard et al., 2015), as well as development (Quintana et al., 2010) and even lifespan in animal models (Akbari et al., 2019; Latorre-Pellicer et al., 2016). But in human studies, we currently lack tools to track changes in mitochondrial biology from non-invasive biofluids such as saliva.

An emerging biomarker of mitochondrial stress and signaling is the circulating mitochondrial genome, which is released as cell-free mitochondrial DNA (cf-mtDNA) (Trumpff et al., 2021). In the blood, measuring cf-mtDNA cross-sectionally (one single collection) showed that elevated cf-mtDNA levels are associated with physical trauma (Lam et al., 2004), psychopathology (Lindqvist et al., 2016; Lindqvist et al., 2018), infectious conditions (Nakahira et al., 2013), and chronological age (Pinti et al., 2014). In the absence of dynamic time-course data, these findings led to the idea that cf-mtDNA is a relatively stable marker of disease. But subsequent studies showed that blood cf-mtDNA is rapidly inducible within 5-30 minutes after psychological stress (Hummel et al., 2018; Trumpff et al., 2019), and that within-person cf-mtDNA levels may change several-fold from week-to-week or month-to-month (Trumpff et al., 2021), thus challenging the notion that cf-mtDNA is a stable state biomarker. Moreover, cf-mtDNA has been detected not only in blood but also in other biofluids, including urine (Kim et al., 2019) and cerebrospinal fluid (Varhaug et al., 2017), indicating that cf-mtDNA may ubiquitously exist in multiple biofluids.

Together, the emerging role of cf-mtDNA in relation to health and disease states, the stress-inducibility and within-person variation in blood cf-mtDNA, and its existence in multiple biofluids, led us to hypothesize that cf-mtDNA may share some dynamic properties with other neuroendocrine factors, such as cortisol, and that it may be detectable in saliva. Here we examine this hypothesis, with a focus on examining the existence and dynamic properties of salivary cf-mtDNA. We also conduct exploratory analyses to examine the association of cf-mtDNA with cortisol, as well as self-reported daily mood and health-related behaviors.

## B. Results

### 1. cf-mtDNA is detectable in saliva and is independent of cellular content

To determine if cf-mtDNA is detectable in saliva collected using standard salivary cortisol procedures (Kirschbaum and Hellhammer, 1994), saliva was collected using salivettes from a small discovery cohort of healthy adults (n=64 samples from 4 women and 3 men, aged 21-72 years) at specific times of the day: at awakening, +30 min, and +45min after awakening. Saliva was extracted using standard procedures for cortisol, followed by an additional centrifugation step (2,000g x 10 min) to eliminate any potential contaminating cells or debris that would influence DNA levels (**Fig S1A**). The resulting cell-free saliva was then lysed to release cell-free DNA (cf-DNA), which was quantified by quantitative PCR (qPCR). To increase the robustness of our cf-DNA quantification, we simultaneously targeted two mitochondrial (mtDNA) and two nuclear (nDNA) amplicons (**Fig S2A**), as in previous work (Trumpff et al., 2019). As previously observed in other human biofluids, we detected substantial amounts of cf-mtDNA in human saliva. Moreover, salivary cf-mtDNA levels were within the physiological range of plasma cf-mtDNA levels detected using the same methodology in an unrelated set of reference control samples (**Fig 1B**).

**Figure 1.**
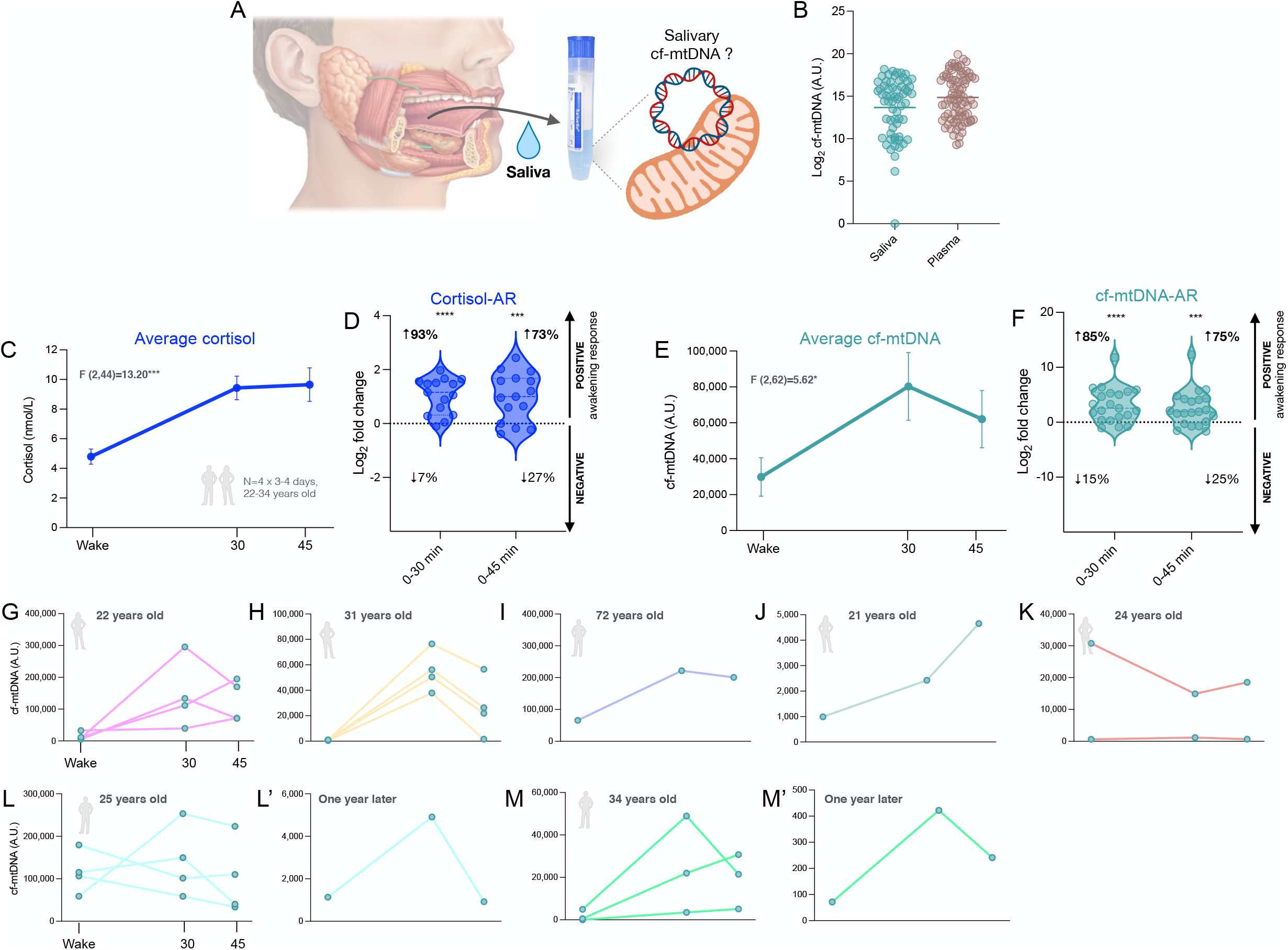
Detectable levels of cf-mtDNA and the awakening response in human saliva. (**A**) The goal of this study was to assess whether cf-mtDNA is detectable in saliva. (**B**) Distribution of cf-mtDNA (lines represent the mean) measured in saliva (n=65 observations in 4 women and 3 men, aged 21-72 years old) and plasma (n=85 observations measured in an independent reference cohort from healthy participants). The data was log(2) transformed prior to visualization. (**C**) To assess potential salivary cf-mtDNA awakening response, saliva was collected at awakening, 30 and 45 min after awakening. Cortisol awakening response in a subset of 4 participants (22 and 31 years old women, 25 and 34 years old men) studied over 3-4 consecutive days. (**D**) Distribution of fold change from awakening to 30 min (left) and awakening to 45 min (right) (15 assessments from a total of 4 participants) in cortisol levels. (**E**) Salivary cf-mtDNA values for 7 participants (n=65 observations in 4 women, 3 men, aged 21-72 years old). Some participants were studied over 3-4 consecutive days and 2 were studied again one year later. (**F**) Distribution of fold change from awakening to 30 min (left) and awakening to 45 min (right) (20 assessments from a total of 7 participants) in cf-mtDNA levels. (**G-M**) Individuals profiles of cf-mtDNA awakening response. Data is shown as means ± SEM. P-values from One-way ANOVA for repeated measures (main effect) (B,D), from chi-squared test comparing proportion of positive and negative awakening responses compared to chance (50:50) (C, E), p<0.05*, p<0.01**, p<0.001***, p<0.0001****.

Saliva contains a large number of buccal epithelial cells and leukocytes (Theda et al., 2018), which could theoretically actively or passively release mtDNA during isolation. To test whether cellular contamination may represent a source of cf-mtDNA, we verified whether cf-mtDNA was influenced by the cellular content of saliva by quantifying the number of cells in each collected saliva sample (**Fig S1B**). The number of cells quantified by qPCR did not correlate with cf-mtDNA levels (r^2^=0.006, N.S.) (**Fig S1C**), thus ruling out cellular content as a substantial source of salivary cf-mtDNA.

### 2. Sedimentation characteristics of salivary cf-mtDNA and collection methods

As an additional test of specificity, we examined the sedimentation characteristics of salivary cf-mtDNA. In blood, cf-mtDNA exists in different forms, including encapsulated within whole mitochondria, within extracellular vesicles, or as free floating DNA (Trumpff et al., 2021). If cf-mtDNA were contained in whole mitochondria, large extracellular vesicles, or apoptotic bodies, it would sediment and be depleted during high speed centrifugation (18,000g x 15 min) (Dache et al., 2020). On the other hand, small vesicles or free-floating naked forms of cf-mtDNA would not. In our saliva samples, high-speed centrifugation decreased cf-mtDNA levels by only 17% on average (small effect sizes, Cohen’s d=0.13-0.18, **Fig S1E**). Thus, less than one-fifth of saliva cf-mtDNA is contained in large vesicular structures (e.g., whole mitochondria or extracellular vesicles) whereas >80% is either naked or contained in very small vesicular structures. These results using the first mtDNA amplicon (ND1) were confirmed with a second mtDNA amplicon (COX1), which showed high concordance (**Fig S2B-D**). Therefore, the remainder of our analyses use the average of both amplicons as a stable estimate of total salivary cf-mtDNA.

For typical salivary hormone analyses, saliva is collected either with salivettes (as above) or by passive drool. To evaluate the impact of the saliva collection method on cf-mtDNA levels, we compared cf-mtDNA and cf-nDNA levels from saliva collected using both methods. Participants collected saliva using both methods in a randomized, cross-over order. We observed that thawed passive drool samples tended to agglutinate, and they contained 93% less cf-mtDNA than salivette-derived saliva (**Fig S3A-B**). cf-mtDNA levels measured by both methods were weakly correlated (r^2^=0.05-0.10). Moreover, compared to passive drool where cf-mtDNA values were narrowly distributed near the lower limit of detection, salivette-derived saliva from the same individuals showed a higher dynamic range, suggesting that the salivette approach is technically superior (**Fig S3C-D**). Finally, we confirmed that freezing salivettes at -20 or -80°C after collection, which is often required for long-term storage in some studies, did not significantly alter cf-mtDNA levels (**Fig S3E-F**). Thus, the standard method of saliva collection, storage, and processing used for cortisol measurements are also appropriate to quantify cf-mtDNA. Having demonstrated that cf-mtDNA is i) abundant in human saliva, ii) independent of cellular content, and iii) robust to storage and handling conditions, provided the basis to examine cf-mtDNA dynamics.

### 3. Saliva cf-mtDNA exhibits an awakening response

To establish whether salivary cf-mtDNA exhibits an awakening response, we collected saliva at three timepoints recommended to quantify the CAR: awakening, +30 and +45 min (Stalder et al., 2016). As expected, we observed relatively consistent CARs across individuals, with cortisol increasing on average from 4.80 to 10.99 mMol from awakening to peak (+129.06%, P<0.0001, n=15 days, from 4 individuals, **Fig 1C**). A positive CAR was observed on 93% (awakening to +30 min) and 73% (awakening to +45 min) of days (**Fig 1D**), confirming participant compliance and the adequacy of the saliva sampling method for cortisol. Awakening responses for other steroid hormones including cortisone, testosterone, progesterone, dehydroepiandrosterone (DHEA), and estradiol are shown in **Fig S4**.

Quantifying cf-mtDNA at the same timepoints revealed that salivary cf-mtDNA similarly exhibited a robust awakening response. On average in this small sample, cf-mtDNA increased by 2.69-fold from 0 to +30 min (p<0.05) and remained elevated at +45 min (**Fig 1E**, individual trajectories are shown in **Fig 1G-M**). The highest cf-mtDNA awakening response (cf-mtDNA-AR) was a 83.5-fold increase (from 915 to 76,525 A.U, 31-year-old woman, **Fig. 1H**). Similar to cortisol, a positive cf-mtDNA-AR was detected on 85% of days, demonstrating a consistent cf-mtDNA-AR similar in dynamics to the CAR (**Fig 1F**). Self-reported compliance with the timing of collection immediately at awakening was associated with the magnitude of the cf-mtDNA-AR.

When comparing morning timepoints cf-mtDNA levels between participants, unlike some other physiological parameters such as blood pressure and glycemia which have relatively narrow ranges in healthy individuals, we noted several fold variation in salivary cf-mtDNA levels between individuals. Moreover, important differences in cf-mtDNA-AR were also observed in samples from the same individuals between two occasions of testing separated by ∼1 year (**Fig 1L, M**), indicating large within-person variation in salivary cf-mtDNA dynamics over time. Because establishing the dynamic within-person changes in salivary cf-mtDNA is paramount to assessing its potential utility, we focus the remaining of our investigation on within-person variation.

### 4. Within-person variation in saliva cf-mtDNA

Understanding how salivary cf-mtDNA varies day-to-day and over time is paramount to establish its usability as a biomarker. Therefore, rather than examining cf-mtDNA in a larger cohort that would likely confirm the above results without providing additional information about the temporal dynamics of within-person variation, we performed an intensive repeated-sampling, multi-week protocol in a highly compliant participant, Participant A (34-year-old male, author M.P.). Using the same timepoints as above (awakening, +30 and +45 min) as well as a bedtime timepoint (bedtime) to examine potential diurnal patterns, we quantified cf-mtDNA in parallel with a salivary steroid hormone panel daily for ∼2 months.

Participant A was studied over 53 consecutive days (n=212 observations, 8 missing time points) (**Fig 2A**). Consistent with the results above and with the literature, a positive CAR was detected on 80% of days (p<0.0001 +30 min, p<0.01 +45min), with an average cortisol increase of 30% (5.6 to 7.6 mMol), and a daytime decline from morning peak to bedtime was observed on 98% of days (p<0.0001, **Fig 2B, D**). The awakening response patterns for other steroid hormones are shown in (**Fig S5**).

**Figure 2.**
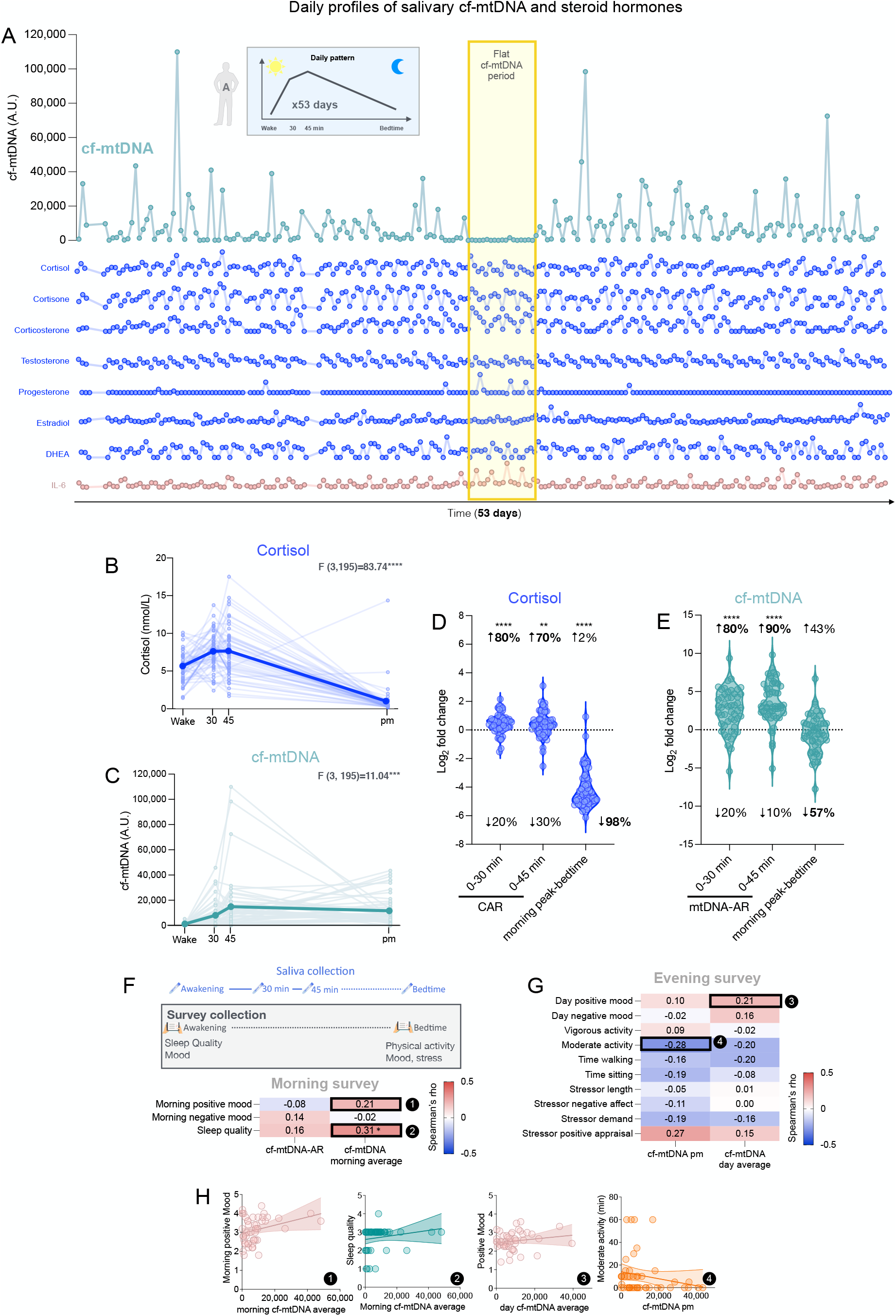
Salivary cf-mtDNA diurnal variation in participant A. (**A**) Day-to-day variation of salivary cf-mtDNA, steroid hormones and IL-6 in a 34 year-old male participant (Participant A). (*Top*) Salivary cf-mtDNA collected at awakening, 30 and 45 min after awakening, and bedtime over 53 consecutive days (n=212 observations). 53-day diurnal profiles for (**B**) cortisol and (**C**) cf-mtDNA. Trajectories for each hormone are shown in Figure S9. (**D**) Fold change in saliva cortisol from awakening to +30, awakening to +45 and morning peak to evening values. (**E**) Same as in (D) for salivary cf-mtDNA. (**F**) Exploratory analyses of the associations between cf-mtDNA levels, mood, and health-related behaviors from the daily diaries collected in the morning and (**G**) evening. Numbers are the effect sizes (Spearman rho), color-coded by the directionality of the association (*blue*, negative; *red*, positive). (**H**) Selected examples of association shown as scatterplots (numbered 1-4) from F and G. In (B, C) data is shown as means and individual datapoints. P-value from One-way ANOVA for repeated measures (main effect). (D, E) P-value from chi-squared test comparing proportion of positive and negative awakening response compared to chance (50:50). (F) P-value from Spearman’s correlation test, p<0.05*, p<0.01**, p<0.001***, p<0.0001***.

Across the 53 days, Participant A exhibited a robust cf-mtDNA-AR, where salivary cf-mtDNA increased on average 17.2-fold from awakening to +45min (P=0.0004) (**Fig 2C, E**). This diurnal cf-mtDNA pattern cannot be driven by cellular contamination because salivary cellular content follows an opposite diurnal pattern; while cf-mtDNA increases after awakening, salivary cell count decreases 2.1-fold from awakening to +30/45min (**Fig S1D**). The maximal detected cf-mtDNA-AR (Day 35) was an 892-fold elevation over the first 45 min after awakening. These observations are in line with the data in Figure 2, albeit of larger overall magnitude, and also of substantially greater magnitude than what is observed for typical hormones such as cortisol. These results also revealed large natural day-to-day variation (112% CV).

To explore potential psychological and behavioral factors associated with day-to-day variation, we collected daily measure of sleep quality (morning), affective states (morning and evening), physical activity (evening), and stressor exposure (evening) (**Fig 2F-H**, individual trajectories are shown in **Fig S6A**). A small positive association was found between sleep quality and average morning cf-mtDNA levels (rho=0.31, p<0.05, not adjusted for multiple testing), but the factors influencing cf-mtDNA variation still remain to be understood.

These high temporal resolution results called for a similar study in another participant. Therefore, a second Participant B (35-year-old male, author G.G.), age- and sex-matched to Participant A, was studied over 60 consecutive days (n=220 observations, 20 missing time points) (**Fig 3A**). A positive CAR was found on 66% of the days (p=0.021 at +30, p=0.045 at +45), with an average magnitude of +90% (6.4 to 12.2 mMol), and a daytime decline from morning peak to bedtime was observed on 100% of the days (p<0.0001) (**Fig 3B, D**), confirming the adequacy of the saliva sampling methodology, and normal HPA axis function. See **Fig S5G-K** for other steroid hormones awakening patterns for Participant B.

**Figure 3.**
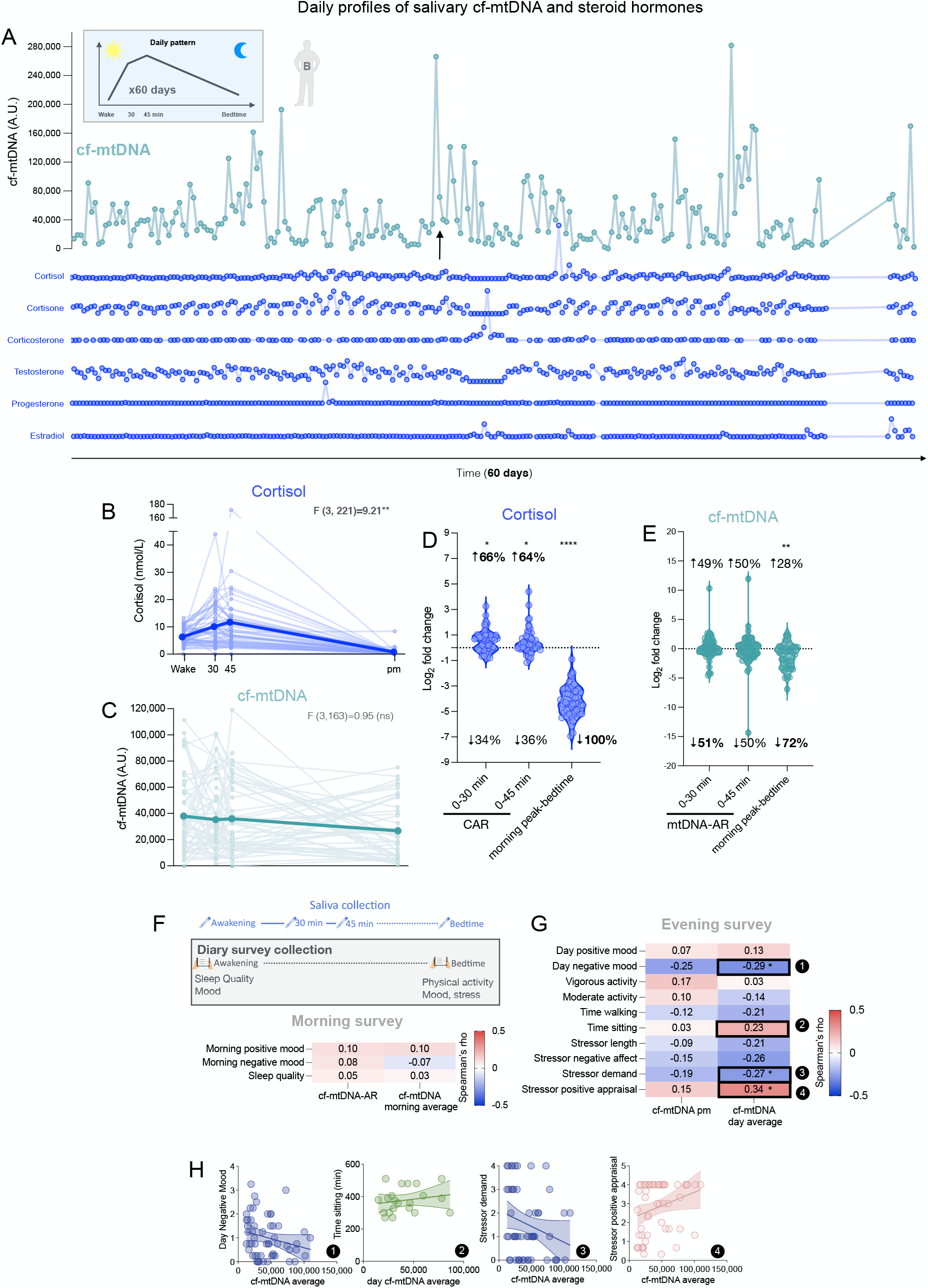
Salivary cf-mtDNA diurnal variation in Participant B. (**A**) Day-to-day variation of salivary cf-mtDNA, steroid hormones in a 35 year-old male participant (Participant B). (*Top*) Salivary cf-mtDNA collected at awakening, 30 and 45 min after awakening, and bedtime over 60 consecutive days (n=220 observations). 60-day diurnal profiles for (**B**) cortisol and (**C**) cf-mtDNA (without outliers). (**D**) Fold change in saliva cortisol from awakening to +30, awakening to +45 and morning peak to evening values. (**E**) Same as in (D) for salivary cf-mtDNA. ((**F**) Exploratory analyses of the associations between cf-mtDNA levels, mood, and health-related behaviors from the daily diaries collected in the morning and (**G**) evening. Numbers are the effect sizes (Spearman rho), color-coded by the directionality of the association (*blue*, negative; *red*, positive). (**H**) Selected examples of association shown as scatterplots (numbered 1-4) from F and G. In (B, C) data is shown as means and individual datapoints. (D, E) P-value from chi-squared test comparing proportion of positive and negative awakening response compared to chance (50:50). (F) P-value from Spearman’s correlation test, p<0.05*, p<0.01**, p<0.001***, p<0.0001***.

In contrast to the normal CAR profile, the average salivary cf-mtDNA-AR pattern in Participant B was not consistent, showing an average 5-8% decline from awakening to +30/45min (**Fig 3E**). Some days exhibited a robust awakening response reaching up to 14.59-fold (**Fig 3C**). Consistent with the cohort and Participant A, a cf-mtDNA morning peak to bedtime decline was observed on 72% of the days (p=0.0003) (**Fig 3E**). Because the cf-mtDNA patterns of participant B differed from the ones of the cohort and of participant A, all Participant B samples were re-assayed, which validated these results and showed high internal validity for the salivary cf-mtDNA assay (r^2^=0.84, p<0.0001, **Fig S7**). In this participant also, there were large (50% CV) day-to-day variation in the average salivary cf-mtDNA levels.

In relation to psychological and health-related behavior factors, Participant B showed higher salivary cf-mtDNA levels on days with higher levels of positive appraisal of daily stressors (rho=0.34, p<0.05, **Fig 3F-H**), individual trajectories are shown in **Fig S6B**). Days with lower cf-mtDNA levels were characterized by greater scores for negative affective states (rho=-0.29, p<0.05) and stressor demand (rho=-0.27, p<0.05). Overall, as for participant A, the daily diary data and self-reported behaviors accounted for a limited portion of the day-to-day variation in cf-mtDNA.

### 5. Associations of saliva cf-mtDNA dynamics and steroid hormones

Performing repeated-measures in the same participants makes possible to identify specific periods of time that deviate substantially from that individual’s baseline or typical pattern (Poldrack et al., 2015), which provides a unique scientific opportunity for discovery not available in cross-sectional designs. In Participant A, we identified a 4-day period (days 27-30) where absolute cf-mtDNA levels were abnormally low (97% lower than normal, p=0.014) and the cf-mtDNA-AR was strikingly absent. On those days, cortisol levels also were abnormal, marked by a complete absence of CAR (**Fig S8**), which was unusual (deviation by 2 S.D. from average behavior, p=0.0057) for this participant.

This finding prompted us to examine the correlation between salivary cf-mtDNA and cortisol. Across the 212 observations for participant A, the cortisol and cf-mtDNA awakening responses were moderately positively correlated (r^2^=0.19, p=0.0015) (**Fig S9A**). While these results were not replicated in participant B (r^2^=0.00, p=0.28), a similar effect size was found for this association in the participant cohort (N=14 days from 4 individuals, r^2^=0.17, p=0.14) (**Fig S9B-C**). Together, this suggested a potential association between cortisol and cf-mtDNA dynamics, consistent with previous work in blood and cultured cells (Lindqvist et al., 2016; Trumpff et al., 2019). Moreover, visual inspection of the cf-mtDNA and steroid hormone patterns revealed that during the flat period when cf-mtDNA levels were abnormally low, progesterone levels were abnormally elevated by 1.38-fold above average (p<0.0001).

To explore the cortisol/cf-mtDNA association in Participant B in a hypothesis-driven manner, we isolated days with a robust CAR (+2-fold or higher from awakening to +30 min, n =18) and days without a CAR (n=12), but did not find a significant difference in the cf-mtDNA-AR between those periods (**Fig S10**). We note that on average, the steroid hormone patterns differed between both repeat participants. Compared to Participant A, Participant B showed a 1.3-fold higher average cortisol levels (p<0.03), and higher daily variability for cortisol (2.6-fold), corticosterone (5-fold), and progesterone (20-fold) (**Fig S11**). During the study period, Participant B underwent a regimented marathon running training program, completed a marathon on Day 27 (arrow in **Fig 3A**), and reported several objective major life stressors. Altogether, this suggests that the two participants may not be directly comparable, highlighting with high confidence the existence of inter-individual variation in salivary steroid hormones and cf-mtDNA dynamics.

### 6. Saliva cf-mtDNA is negatively associated with IL-6 saliva levels at bedtime

Finally, given the mixed evidence in the literature linking cf-mtDNA and pro-inflammatory states (Trumpff et al., 2021), we again leveraged the high-frequency, repeated-measures saliva measurements in Participant A to examine if the salivary cf-mtDNA-AR was associated with the pro-inflammatory cytokine IL-6. Consistent with prior literature in plasma/serum (Nilsonne et al., 2016) and saliva (Izawa et al., 2013), salivary IL-6 exhibited a strong negative awakening response (**Fig 4A**). IL-6 levels were highest at awakening, decreased on average by 85-88% over the first 30-45min (P<0.0001), and increased by an average of 4.0-fold from +45 min to bedtime (P<0.0001). These results confirm that IL-6 and cf-mtDNA exhibit opposite awakening responses (**Fig 4B)**.

**Figure 4.**
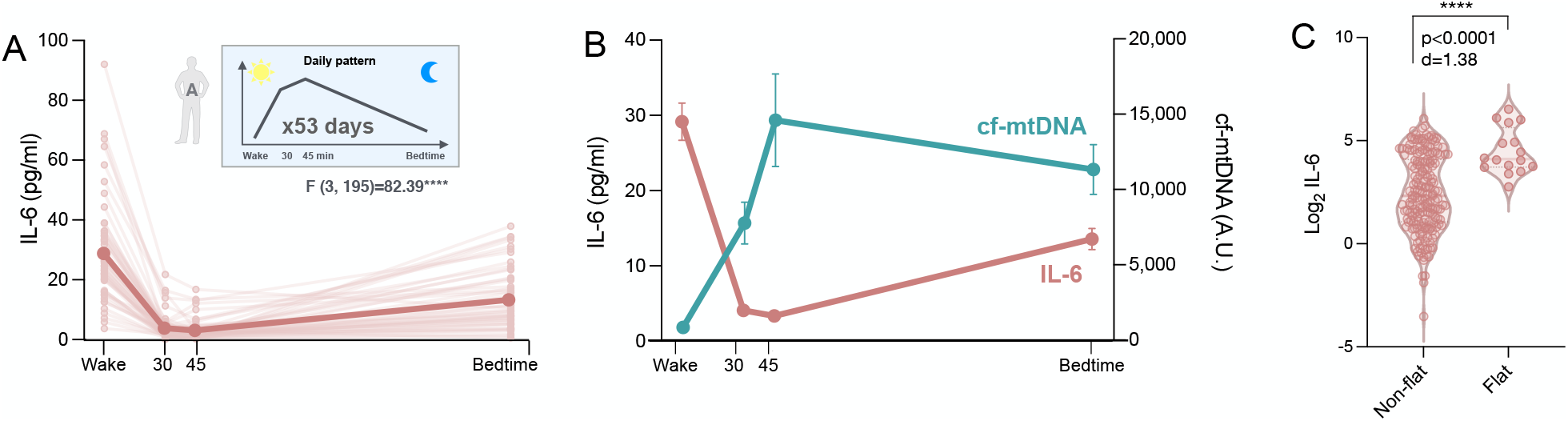
Salivary IL-6 and cf-mtDNA diurnal variation. (**A**) IL-6 diurnal patterns in participant A with data collected over 53 days at awakening, +30 min, +45 min and evening (n=204 observations). Data is shown as means and individual datapoints. P-value from One-way ANOVA for repeated measures (main effect). (**B**) Trajectories of salivary IL-6 and cf-mtDNA. Data is shown as means ± SEM. (**C**) Comparison of IL-6 levels on normal days (non-flat) and days with unusually low cf-mtDNA levels (flat) (see figure S7). IL-6 values were log2 transformed prior to visualization. P-value from unpaired t-test. p<0.0001****.

The opposite patterns of change between salivary cf-mtDNA and IL-6 over time leads to two conclusions. First, this provides additional evidence for the specificity of the cf-mtDNA dynamics, demonstrating that the observed changes over time are not driven by the dilution or concentration of saliva (which would change the concentration of all analytes in the same direction and in a similar magnitude). This point was further confirmed by quantifying total salivary protein concentration at multiple time points across multiple consecutive days. Saliva concentration was relatively stable and did not vary systematically across the day, and it was not correlated with cf-mtDNA (**Fig S12**). Second, salivary IL-6 and cf-mtDNA are modestly but significantly negatively associated (r^2^ = 0.08, p<0.0001, **Fig S13A**), particularly at bedtime (r^2^ = 0.11, p = 0.016, **Fig S13B**). Moreover, compared to the 49 days with normal cf-mtDNA-ARs in Participant A, the flat 4-day period with abnormally low cf-mtDNA exhibited markedly elevated IL-6 levels (2.75-fold, p<0.0001) (**Fig 4C**). This result corroborates the negative association between cf-mtDNA and the pro-inflammatory cytokine IL-6 (see **Fig S13A**).

## C. Discussion

We have reported the existence and dynamic behavior of cf-mtDNA in human saliva. Similar to other neuroendocrine factors that exhibit natural diurnal variation and a robust awakening response, such as cortisol, our results show that salivary cf-mtDNA exhibits a large magnitude awakening response (cf-mtDNA-AR). The high-frequency, daily repeated-measures strategy across several weeks also demonstrated high day-to-day variation in salivary cf-mtDNA, as well as relatively large stable inter-individual differences between both participants. Together, these data open the door to using salivary cf-mtDNA as a scalable, minimally invasive outcome measure to capture both inter-individual differences, and also dynamic intra-individual variation in some aspect of mitochondrial biology and signaling in relation to health-related outcomes.

Mitochondria are not only the major source of cellular energy, but also contribute to the biosynthesis of several neuroendocrine factors. Cortisol, as well as all steroid hormones – mineralocorticoids, estrogens, androgens – are in fact enzymatically produced within the mitochondrial matrix (Midzak and Papadopoulos, 2016). The first and last enzymatic reactions that produce cortisol take place inside mitochondria of the adrenal gland zona fasciculata. For testosterone the first step takes place in Leydig cells mitochondria within the testes, and for estrogens and progesterone mainly within the ovaries’ mitochondria (Midzak and Papadopoulos, 2016). Naturally, the circulating mitochondrial genome (i.e., cf-mtDNA) is also derived from mitochondria, although its exact tissue(s) or cell(s) of origin remain unclear. Other “uniquely mitochondrial” proteins such as the heat shock protein 60 (HSP60) are released systemically in the blood. In population-based and clinical studies, mitochondrial HSP60 was elevated in individuals with higher levels of psychosocial distress (Shamaei-Tousi et al., 2007) and with cardiovascular disease (Pockley et al., 2000). In our data, the generally small correlations found between cortisol and cf-mtDNA make it unlikely that cortisol and cf-mtDNA would be co-released from the same mitochondria. Rather, we speculate that distinct mitochondria-derived signals are released by different source mitochondria (adrenal glands mitochondria for cortisol, other tissue(s) for cf-mtDNA) to fulfill different regulatory roles within the organism (Picard et al., 2018). The physiological role of salivary cf-mtDNA is completely unknown.

Another major knowledge gap relates to the mechanism responsible for the rapid release and dynamics of cf-mtDNA. Studies in blood showed that cf-mtDNA can change by 2-3-fold in serum and plasma over 5-30 minutes (Hummel et al., 2018; Trumpff et al., 2019), and that cf-mtDNA levels measured at one timepoint can vary up to an order of magnitude when measured again within weeks or months (Trumpff et al., 2021). These data suggested that blood cf-mtDNA level is not a stable trait of the individual, but may have a dynamic behavior that reflect changeable states (Trumpff et al., 2021). Our saliva data corroborate this assertion by showing that salivary cf-mtDNA can increase by over 100-fold over periods of 15-30 minutes, establishing its highly dynamic nature. In comparison, salivary cortisol increases by 2-to-5 fold within 20-30 minutes post-stress, or more moderately after awakening, and decreases with similar kinetics (Foley and Kirschbaum, 2010). Based on these data, cf-mtDNA and cortisol appear to have similar kinetics but differ mainly in the magnitude of their variation. Higher temporal resolution data are necessary to fully define its temporal dynamics. In general, our measurements show that the salivary cf-mtDNA-AR is likely ∼10-100 fold larger than the average salivary CAR. Mechanistically, cortisol and cf-mtDNA were only weakly correlated, suggesting they are either unlikely causally linked to one another, or that cortisol in itself may contribute but is not sufficient to increase salivary cf-mtDNA.

We also completed exploratory micro-investigations into the associations between psychological and behavioral factors with cf-mtDNA for each of our highly studied participants (Participants A and B). These analyses revealed inconsistent and inconclusive effects within each of these participants. However, these effects do not preclude considering testing variability among psychological and behavioral factors, within and between participants, together with day-to-day and between person fluctuations in cf-mtDNA. Using a simultaneous within- and between-subject study design would be useful to understand the individual determinants of salivary (and blood) cf-mtDNA levels, as others have done with cortisol (Adam et al., 2006).

Technically, our confidence that cf-mtDNA is a robust and specific biological entity is enhanced by three observations. First, cf-mtDNA is independent of the number of cells in saliva, which makes it unlikely that salivary cf-mtDNA is mainly released locally by buccal epithelial cells or leukocytes. This also confirms that cf-mtDNA is not technically confounded by cellular content or other factors related to sample preparation, such as the collection method. Second, the vast majority of cf-mtDNA at different time points exists in non-sedimentable form (i.e., not precipitated by high centrifugal force). This indicates that cf-mtDNA exists in a biological form of transport that is either membrane free, such as naked DNA, or encapsulated by small extracellular vesicles. This observation was consistent across all timepoints analyzed. Third, analyses of total protein concentration and IL-6 concentrations both confirm that the large observed variations in cf-mtDNA cannot be attributed to changes in overall saliva concentration. Instead, this implicates selective cf-mtDNA release mechanisms, as we and others have observed in blood (Dache et al., 2020; Hummel et al., 2018; Trumpff et al., 2019). More research is required to determine the exact form of transport and origin of salivary cf-mtDNA (Trumpff et al., 2021) and whether there is only one major form of salivary cf-mtDNA, or multiple forms. Further research is also needed to establish to what extent salivary cf-mtDNA levels reflect cf-mtDNA in the blood and other biofluids.

Methodologically, we provide an extraction-free, lysis method to study cf-mtDNA levels and dynamics in saliva. The use of saliva has several advantages over more invasive measures in blood: i) saliva samples can be easily collected to study cf-mtDNA behavior in vulnerable populations including children (Ridout et al., 2019), ii) compared to blood or cerebrospinal fluid, saliva collection imposes minimal to no psychological stress/pain, which could influence cf-mtDNA levels (Hummel et al., 2018; Trumpff et al., 2019), and iii) cf-mtDNA measurements (after an additional centrifugation step to eliminate potential cellular contamination) can be performed on stored saliva samples collected with the salivette method. This makes possible salivary cf-mtDNA studies from existing biobanks. Moreover, since salivary cf-mtDNA is robust to freezing effects, saliva samples in new studies can be conveniently collected and stored frozen at home, which can greatly enhance the ecological validity of future mitochondrial psychobiology studies. These factors should facilitate filling the current knowledge gaps around the psychobiological determinants of salivary cf-mtDNA dynamics, and the physiological and health significance of salivary cf-mtDNA from prospective studies.

In summary, our findings demonstrate the existence of cf-mtDNA in human saliva and provide a method to quantify its dynamic behavior, including its robust awakening response and diurnal variation. This finding expands our knowledge of cf-mtDNA in humans to saliva, and defines a broad range of salivary dynamics whose causes and consequences largely remain unexplained. Thus, this work provides a basis for larger well-controlled studies required to understand the significance of salivary cf-mtDNA in relation to health-related factors, including how behaviors and psychosocial stress, as well as changing mental and physical health states, interact with mitochondria.

## D. Method

### Participants

All procedures for this study were approved by New York State Psychiatric Institute (Protocol # 7748). All participants provided written informed consent for their participation and publication of the results. Healthy adults between 18 and 100 years were eligible for inclusion. Exclusion criteria included current diagnosis of cancer or chronic inflammatory disorders, involvement in other clinical trials, and pregnancy. Participants were recruited via flyers posted in the Columbia University Irving Medical Center (CUIMC) community and using a snowball recruitment strategy. The discovery cohort included 7 individuals (4 women, 3 men), aged 21-72 years old. To circumvent the high participant burden and potential cost associated with collecting multiple daily samples across multiple consecutive weeks, repeated daily measures were collected across 53 and 60 days from two healthy Caucasian men (authors M.P., 34 years old, repeat participant A, and G.G., 35 years old, repeat participant B) to assess within-person variability of salivary cf-mtDNA.

### Saliva collection

Saliva was collected at four time points: in the mornings i) immediately upon awakening, ii) 30 minutes after waking up, iii) 45 minutes after waking up, and iv) at bedtime. A total of 1-2 ml of saliva was collected at each sample timepoint using a Starstedt Salivette (Cat# 51.1534.500) following the recommended procedure from the Biomarker Network (Kirschbaum and Hellhammer, 1994). Participants were instructed to place the cotton swab in their mouth (middle of the tongue) for 2-5 minutes, not bite down on the cotton roll, and to avoid contact with the cheeks. Participants then placed the cotton swab back into the Salivette tube and immediately stored the sample in the freezer to preserve analytes (e.g., cf-mtDNA, steroid hormones, IL-6). In the morning, participants avoided brushing their teeth until after the third (+45 min) sample, and the night time sample was collected prior to teeth brushing at bedtime. Participants were instructed not to eat before the +45 min timepoint, and were instructed not to drink water or other fluids 10 minutes before each sample collection. All participants stored salivettes in their freezer (−20°C) before either transporting or shipping samples to the laboratory.

### Salivette processing and storage

Following standard recommended procedures for cortisol, salivettes were thawed and subsequently centrifuged at 1,000 x g for 5 minutes in a refrigerated 4°C centrifuge. From the top fraction of saliva, 500 ul was collected and stored at -80°C for steroid hormone analysis. 200 ul was transferred to a 1.5 ml Eppendorf tube and stored at -80°C for cf-DNA analysis. The remaining saliva was removed, leaving behind ∼50 ul to resuspend total salivary cells (leukocytes and buccal epithelial cells) sedimented at the bottom of the salivette. Cells were collected and transferred to a fresh tube, centrifuged at 2,000 x g for 2 minutes at 4°C, and the supernatant carefully aspirated, leaving behind a clean cell pellet. This cell pellet was used to quantify the total salivary cellular content using the nuclear DNA amplicon B2M.

### Quantification of cf-mtDNA

To ensure the absence of cell and cell debris, each saliva sample was further centrifuged at 2,000 x g for 10 minutes at 4°C prior to processing. To avoid bias in mtDNA content that may occur during DNA isolation using column-based methods (Ware et al., 2020), DNA was extracted using an isolation-free lysis method, adapted from (Picard et al., 2012). 20 ul of saliva was added to 180ul of lysis buffer (500mM Tris HCL, 1% Tween 20, dH_2_0, and 20ug/ml proteinase K), for a dilution of 1:10. Lysis was performed for 10 hours at 55°C, followed by inactivation at 95°C for 10 minutes. This lysis-based method was adapted from a method originally developed for single-cell analysis (Zhou et al., 1997), which allows the detection of small amounts of genomic material without DNA isolation bias inherent to standard column-based DNA isolation techniques (Bender et al., 2006; Grünewald et al., 2016).

Eight ul of the lysed saliva was used directly as template DNA for qPCR. qPCR reactions were run in triplicates using a liquid handling station (ep-Motion5073, Eppendorf) in 384 well qPCR plates. Duplex qPCR reactions with Taqman chemistry were used to simultaneously quantify mtDNA and nDNA amplicons in the same reactions. *Master Mix*_*1*_ for ND1 (mtDNA) and B2M (nDNA) included: Taqman Universal Master mix fast (life technologies #4444964), 300 nM of primers and 100 nM probe (ND1-Fwd: GAGCGATGGTGAGAGCTAAGGT, ND1-Rev: CCCTAAAACCCGCCACATCT, Probe:HEX-CCATCACCCTCTACATCACCGCCC-3IABkFQ.B2M-Fwd:CCAGCAGAGAA TGGAAAGTCAA,B2M-Rev: TCTCTCTCCATTCTTCAGTAAGTCAACT, Probe:FAM-ATGTGTCTGGGTTTCATCCATCCGACA-3IABkFQ). *Master Mix*_*2*_ for COX1 (mtDNA) and RnaseP (nDNA) included: TaqMan Universal Master Mix fast, 300 nM of primers and 100 nM probe (COX1-Fwd: CTAGCAGGTGTCTCCTCTATCT, GAGAAGTAGGACTGCTGTGATTAG, Probe: HEX-TGCCATAACCCAATACCAAACGCC-3IABkFQ. AGATTTGGACCTGCGAGCG, RnaseP-Rev: GAGCGGCTGTCTCCACAAGT, Probe: FAM-TTCTGACCTGAAGGCTCTGCGCG-3IABkFQ. The samples were then cycled in a QuantStudio 7 flex qPCR instrument (Applied Biosystems Cat# 4485701) at 50°C for 2 min, 95°C for 20 sec, 95°C for 1 min, 60°C for 20 sec, for 40 cycles. Reaction volumes were 20 ul. To ensure comparable Ct values across plates and assays, thresholds for fluorescence detection for both mitochondrial and nuclear amplicons were set to 0.08.

cf-DNA was calculated by linearizing the log-based Ct values (2^Ct^) into linear space to obtain quantitative estimates of cf-mtDNA levels in arbitrary units. Because higher cf-mtDNA content is represented by lower Ct values (more template mtDNA copies amplified earlier), the inverse of the linear values were obtained and divided by an arbitrary threshold (1^-12^) that represents the lower detection limit for mtDNA on our instrument (corresponding Ct ∼36). Undetectable cf-DNA values were replaced by 1. The final cf-mtDNA values were obtained from the average of two mtDNA genes (ND1 and COX1) and cf-nDNA was obtained from the average of two single-copy nDNA genes (B2M and RNaseP).

### Steroid hormones

Salivary steroid hormones, including cortisol, cortisone, testosterone, progesterone, corticosterone and dehydroepiandrosterone (DHEA), were quantified using a high throughput liquid chromatography-tandem mass spectrometry (LC-MS/MS) assay as described previously (Gao et al., 2015). The lowest detectable limit was 0.0055 nmol/l for cortisol, 0.0019 ng/ml for cortisone, 2 pg/ml for testosterone, 2.3 pg/ml for progesterone, 5.35 pg/ml for DHEA and 0.9 pg/ml for estradiol. Values below the limit of detection were imputed as the value of the detection limit.

### Salivary protein concentration

Total salivary protein concentration was measured in 44 consecutive samples using the bicinchoninic acid (BCA) protein assay (Bio-World, Cat. #20831001-1), in a 96-well microplate, using endpoint absorbance (optical density, 562 nm OD) on a SpectraMax M3 (Molecular Devices). Saliva samples were diluted 1:10 with nuclease-free water to ensure that all samples were within the assay’s dynamic range, and measured in duplicates. Absolute protein concentration in each sample was determined in a single batch from a standard curve with bovine serum albumin (BSA), adjusting for the dilution factor. Final values are expressed in mg protein per ml of saliva, and ranged from 0.8-4.8mg/ml (median = 1.6 mg/ml).

### IL-6 quantification

Salivary IL-6 concentration was measured by ELISA using the Human IL-6 Quantikine HS ELISA kit (R&D Systems, Cat# HS600C) following the manufacturer’s instructions. After an optimization step, all saliva samples were diluted 1:4 with the diluent, ensuring that all samples were within the assay’s dynamic range. The sample absorbance was determined in microplates on a SpectraMax M3 (Molecular Devices) at 450 nm and 570 nm. A standard curve was used to extrapolate unknown saliva IL-6, multiplied by the dilution factor. Batch-to-batch variations were mathematically corrected using a reference serum sample with known IL-6 concentration included systematically on each IL-6 array. The final IL-6 concentrations are expressed in pg/ml, with a range of 0.1-92 pg/ml (median = 6 pg/ml).

### Diary data

Repeat participants A and B collected measures of mood, stress, sleep quality and physical activity on a daily basis (morning and/or evening) for 53 and 60 days, respectively. The morning diary was completed on awakening and assessed sleep quality using items adapted from the Pittsburgh Sleep Quality Index (PSQI) (Participants were asked to rate “How would you rate the quality of your sleep last night?” on a scale from “very bad” to “very good”)_(Buysse et al., 1989), positive (e.g. Participants were asked to which extent they feel “joyful, glad, happy”, on a scale from “not at all” to “extremely”) and negative (e.g. Participants were asked to which extent they feel “stressed, anxious, overwhelmed”, on a scale from “not at all” to “extremely”) affective states using items adapted from the modified Differential Emotions Scale (MDES) (Fredrickson et al., 2003) as well as expectations on the day to come (e.g. “Please indicate by placing the slider on the line below to what extent you feel the events of your upcoming day are predictable (you know exactly what’s going to happen today)”). The nightly diary was completed at bedtime and assessed mood, daily occurrence of stressors, and physical activity. The intensity of positive (e.g., joy, inspired, feeling in control, etc.) and negative (e.g., feeling stressed, sad, tired, etc.) affective states was measured using items from the MDES (Fredrickson et al., 2003). Stressor occurrence, content (e.g. housing, work, money), duration, and emotional appraisal (e.g. How anxious did you feel at the peak of this stressor? Not at all - extremely) was assessed using items from the daily inventory of stressful events (DISE) (Almeida et al., 2002; Berry Mendes et al., 2007). Finally, physical activity measures included time (in minutes) spent walking and doing moderate, or vigorous physical activities as well as time spent sitting. Summary scores of positive and negative affective states (Fredrickson et al., 2003), positive and negative affective stressors reactivity(Almeida et al., 2002; Berry Mendes et al., 2007) were computed by averaging a subset of items (see supplementary file “Manuscript data” for the list of items included in each scores) and used for the correlation analyses with cf-mtDNA levels.

### Statistical analysis

For both salivary cortisol and cf-mtDNA, Chi-square tests were used to compare proportions of positive and negative awakening response, and proportions of positive and negative slope from morning peak values to bedtime in levels compared to proportions expected by chance (50:50). Bilateral unpaired t-tests were used to compare cf-mtDNA levels obtained using passive drool and salivette, as well as samples frozen and refrigerated. Variation in cf-mtDNA levels and steroid hormones during the day was assessed using repeated measures One way ANOVA. Spearman’s r (r) and/or simple linear regression were used to assess the strength of the associations between cf-mtDNA, diary measures, steroid hormones, and IL-6. Between- and within-person variations were characterized using coefficients of variation (C.V.). DHEA in participant B had more than 50% missing values, so DHEA values were excluded from the analysis. To characterize typical diurnal response, values above 1.5 × (IQR) to the third quartile were removed (in participant B). Statistical analyses were performed with Prism 8 (GraphPad, CA), R version 4.0.2 and RStudio version 1.3.1056. Statistical significance was set at p<0.05.

## Acknowledgements

We would like to acknowledge Celina Porcaro for her work on an initial phase of this project, and Brett Kaufman for useful comments on these data.

## Data availability

All data is available in the supplemental document “Manuscript_data.xlsx”.

