## Supplemental figures for "Dynamic behavior of cell-free mitochondrial DNA in human saliva"

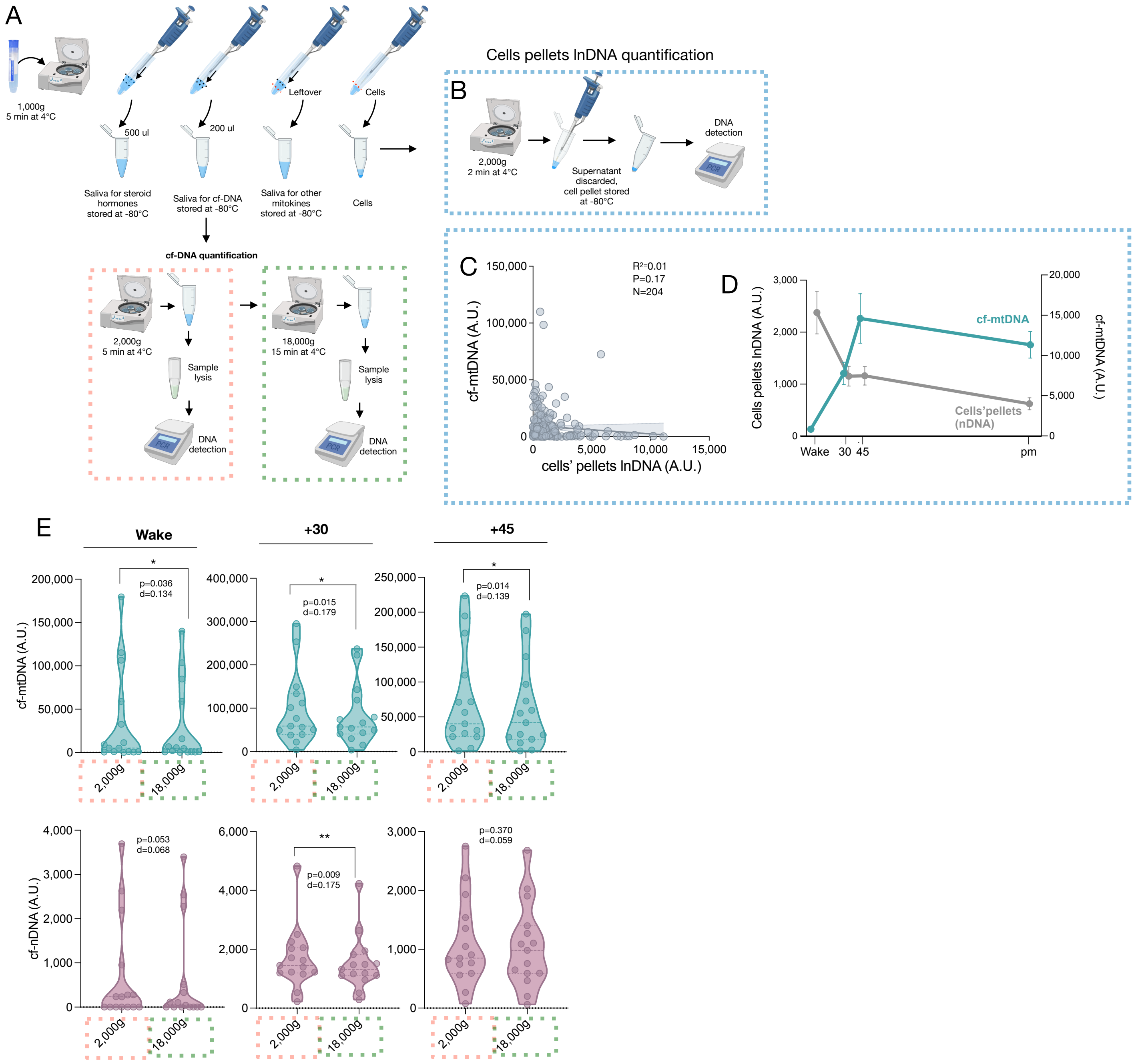


**Figure S1. Characteristics of saliva cf-DNA.** (**A**) Study saliva processing and cf-DNA detection overview. (**B**) Cells' pellet linearized (lnDNA) detection. (**C**) Correlation between salivary cf-mtDNA in participant A and lnDNA measured in saliva's cells pellets (N=212 timepoints). (**D**) Day-to-day variation of salivary cells pellets and cf-mtDNA. Data is shown as means ± SEM. (**E**) Effect of centrifuge speed on cf-DNA in saliva samples collected at awakening, +30min, and +45min, (top, left to right, cf-mtDNA) in 4 participants (22 and 31 year old women 25 and 34 year old men) on 3-4 consecutive days. Paired T test, (bottom, middle, cf-nDNA) Paired T test, *p<0.05, **p<0.01.

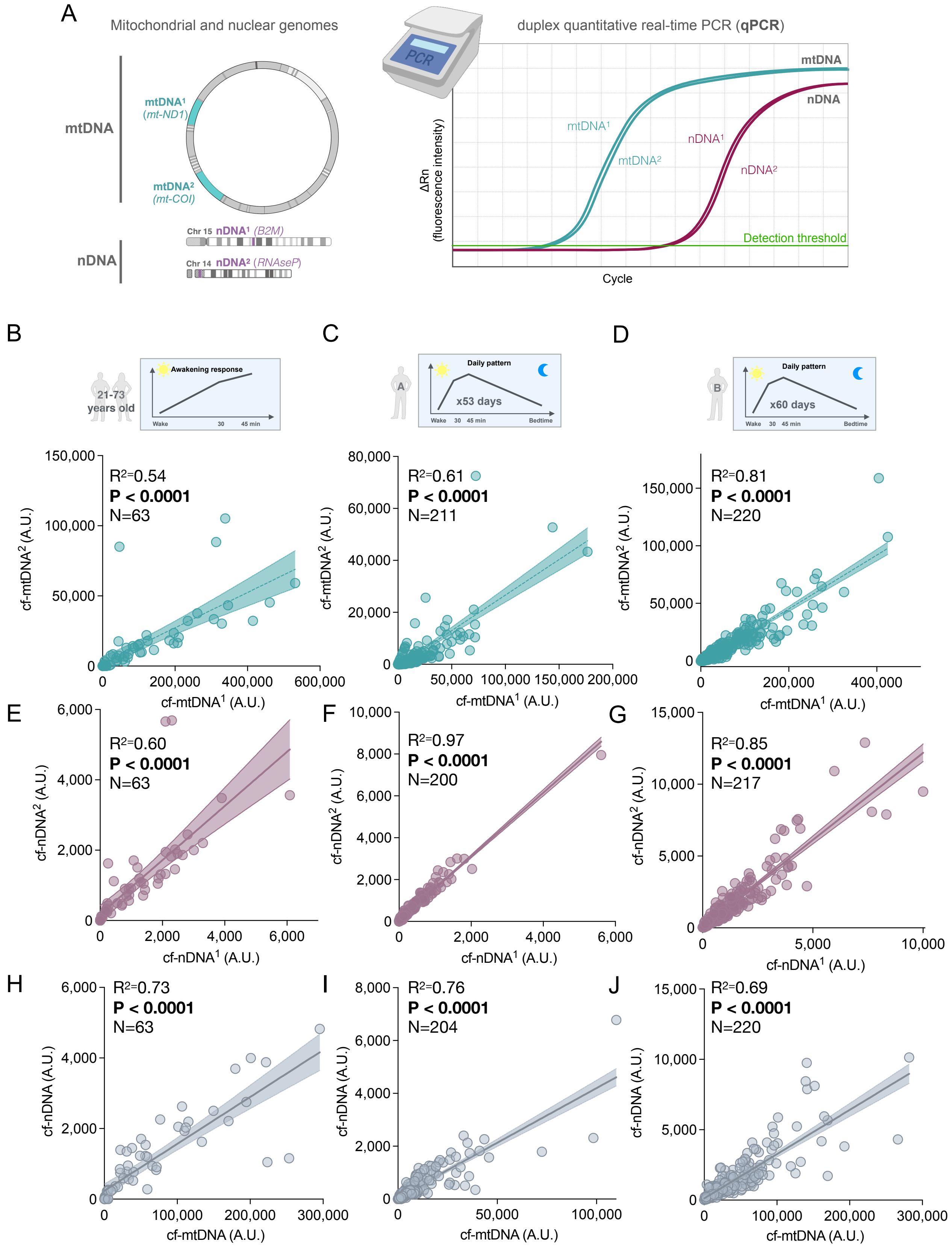


**Figure S2. cf-mtDNA and cf-nDNA detection method.** (**A**) cf-DNA detection method. Correlation between salivary cf-mtDNA amplicons 1 and 2 in samples from (**B**) 7 participants (n=65 observations in 4 women, 3 men, aged 21-72 years old), (**C**) participant A (N=212) and (**D**) B (N=220). (**E-F**) Same as in (B-C) for the correlation between saliva cf-nDNA amplicons 1 and 2. (**H-J**) Same as in (B-C) for the correlation between saliva cf-mtDNA and cf-nDNA (average of amplicons 1 and 2). P-value from simple linear regression.

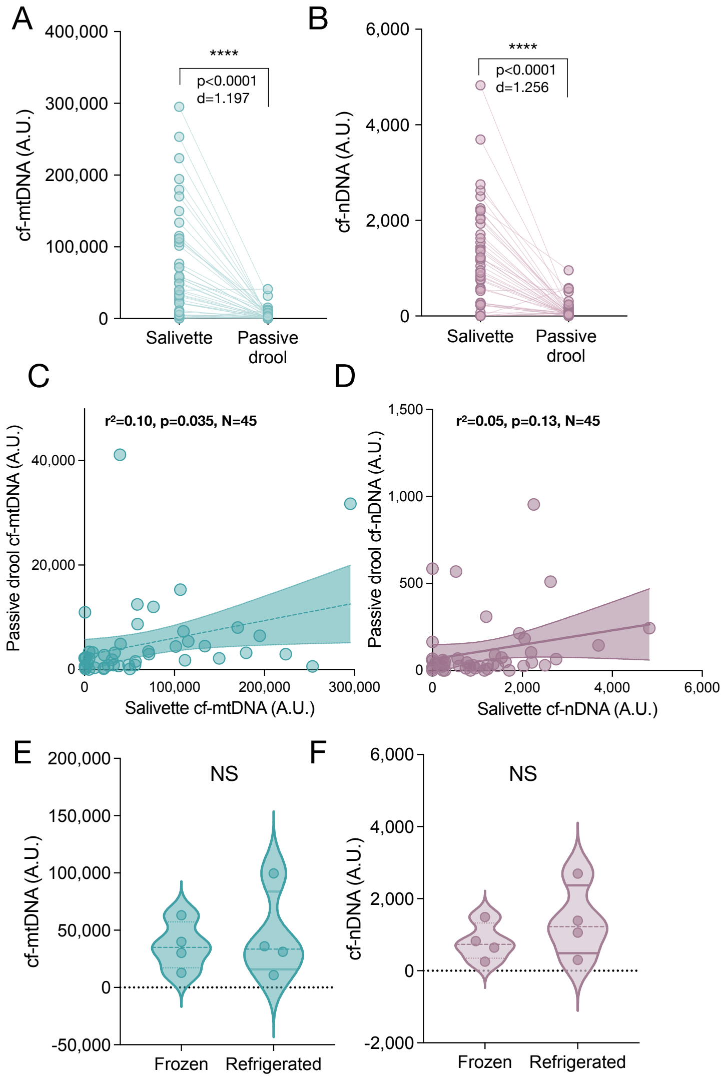


**Figure S3. Effect of collection methods on cf-DNA levels.** Difference between salivette and passive drool collection method on (**A**) cf-mtDNA and (**B**) cf-nDNA levels. Data from n=45 samples from 4 participants at 9-12 timepoints. Correlation between salivette and passive drool collection method in (**C**) cf-mtDNA and (**D**) cf-nDNA levels detected. Effect of saliva storage method (frozen vs refrigerated) on (**E**) cf-mtDNA and (**F**) cf-DNA levels. Unpaired t-test, ****P<0.0001.

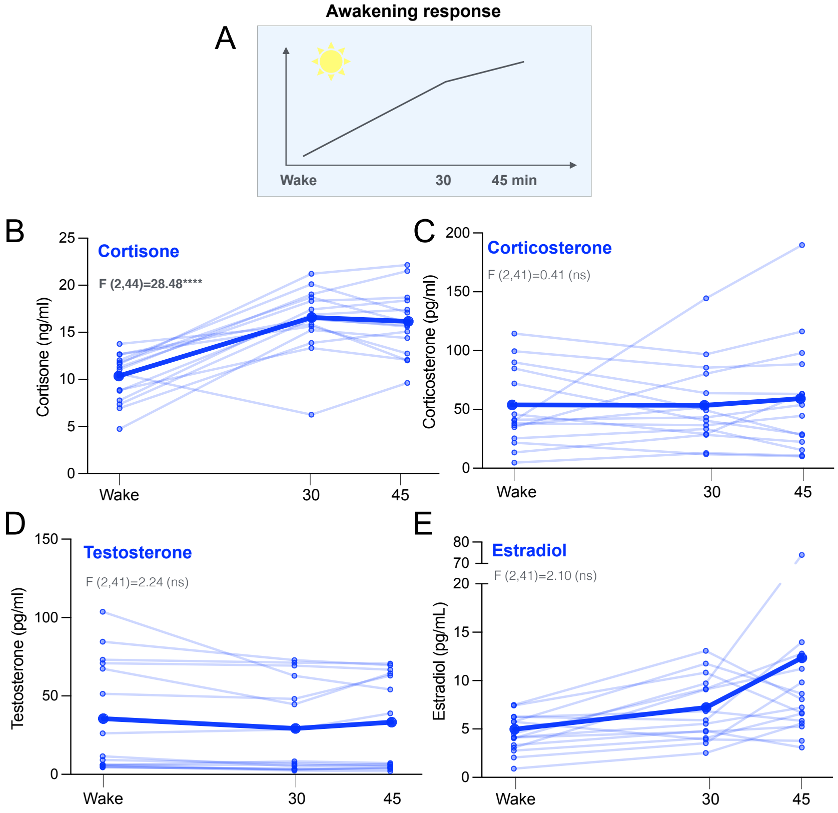


**Figure S4. Steroid hormones awakening response.** (**A**) To assess potential salivary steroid hormones awakening response, saliva was collected at awakening, 30 min and 45 min after awakening in 4 participants (22 and 31 year old women, 25 and 34 year old men). Awakening response for (**B**) cortisone (**C**) corticosterone (**D**) testosterone and (**E**) estradiol. Data is shown as means and individual datapoints. P-value from One-way ANOVA for repeated measures (main effect), p<0.05*, p<0.01**, p<0.001***, p<0.0001***.

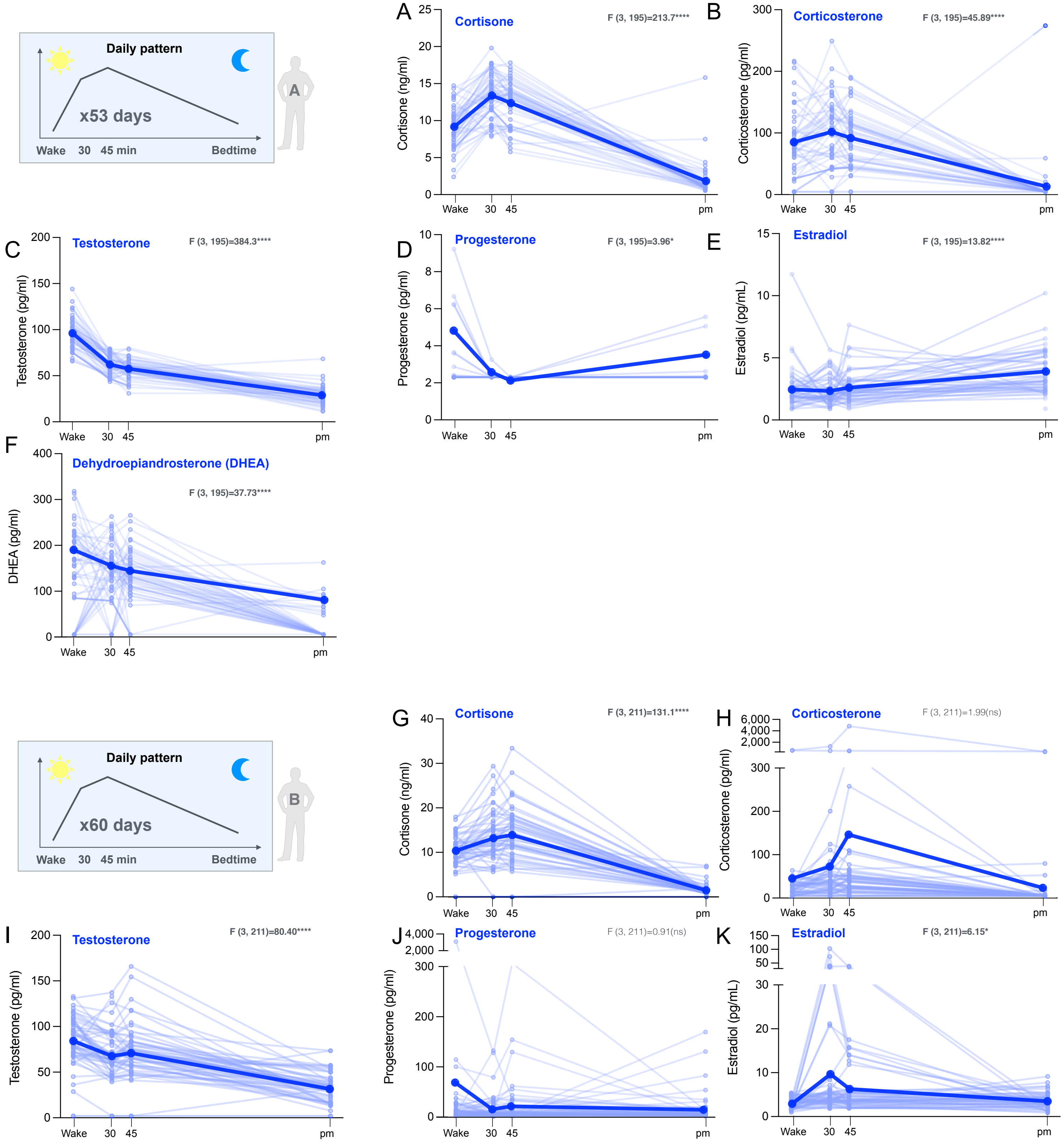


**Figure S5. Steroid hormones awakening response.** Salivary steroid hormones concentration was determined at awakening, 30 and 45 min after awakening and before bedtime in participant A. Awakening response for (**A**) cortisone, (**B**) corticosterone, (**C**) testosterone, (**D**) progesterone, (**E**) estradiol and (**F**) dehydroepiandrosterone (DHEA). (**G-K**) Same as in (A-E) for participant B. DHEA were non-detectable/missing for more than 50% of the timepoints and not analyzed. Data is shown as means and individual datapoints. P-value from One-way ANOVA for repeated measures (main effect), p<0.05*, p<0.01**, p<0.001***, p<0.0001***.

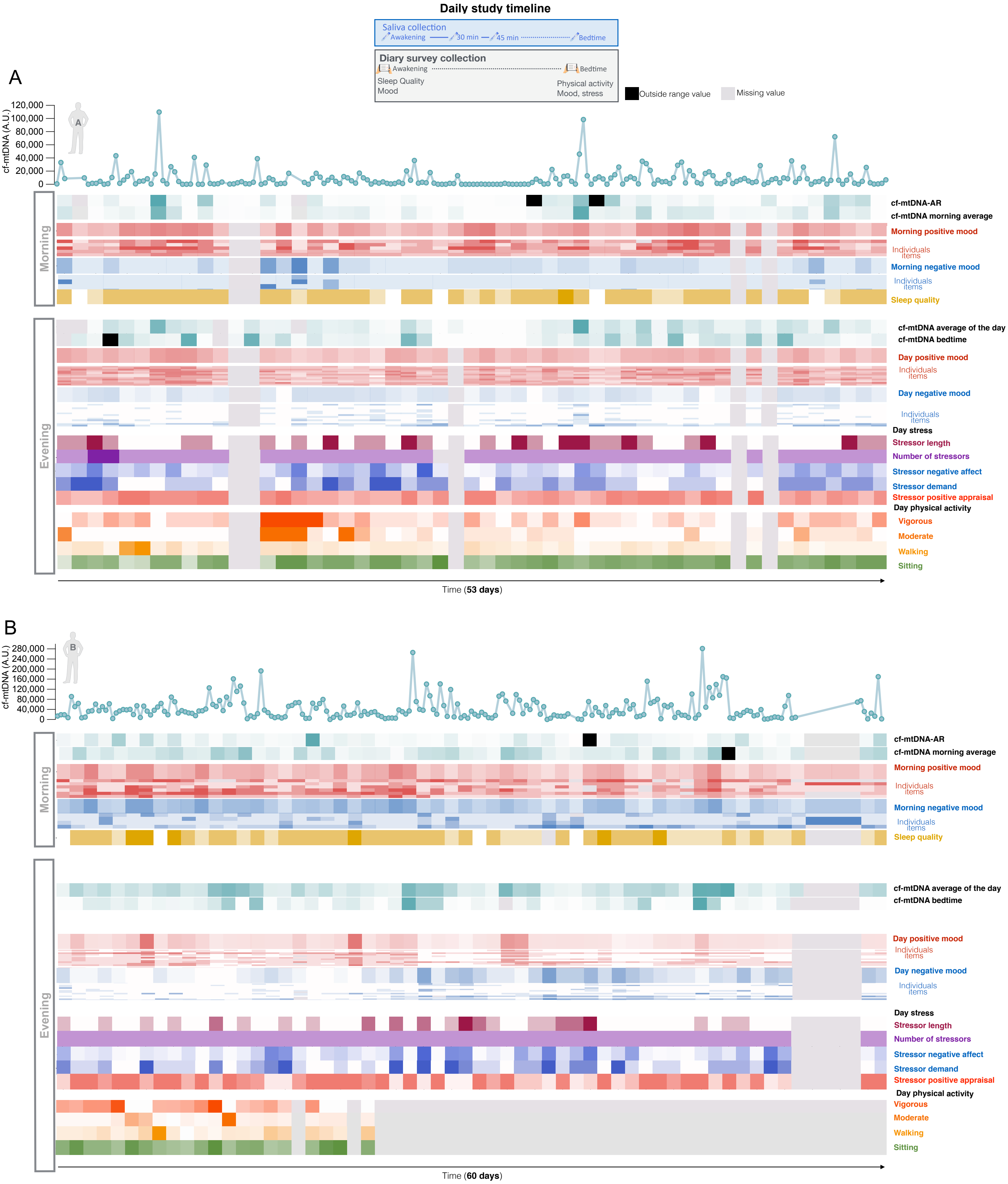


**Figure S6. Salivary cf-mtDNA, affective states, stressors, sleep quality and physical activity diurnal variation.** Heatmaps showing day-to-day variation of salivary cf-mtDNA, mood, sleep quality, stress appraisal and physical activity in (**A**) participant A and (**B**) B.

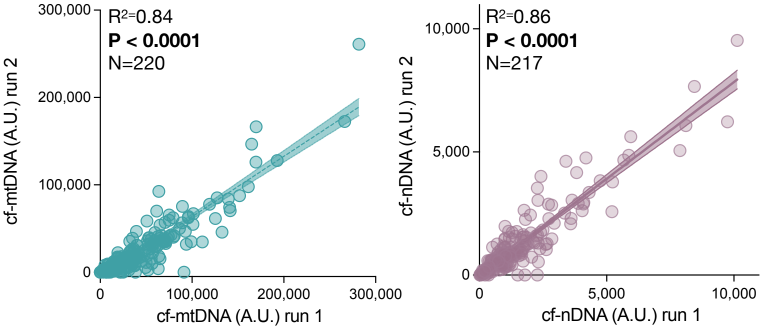


**Figure S7. Saliva cf-mtDNA measured in two different runs.** Comparison of the first and second qPCR runs for (**A**) cf-mtDNA and (**B**) cf-nDNA values in participant B. P-value from simple linear regression.

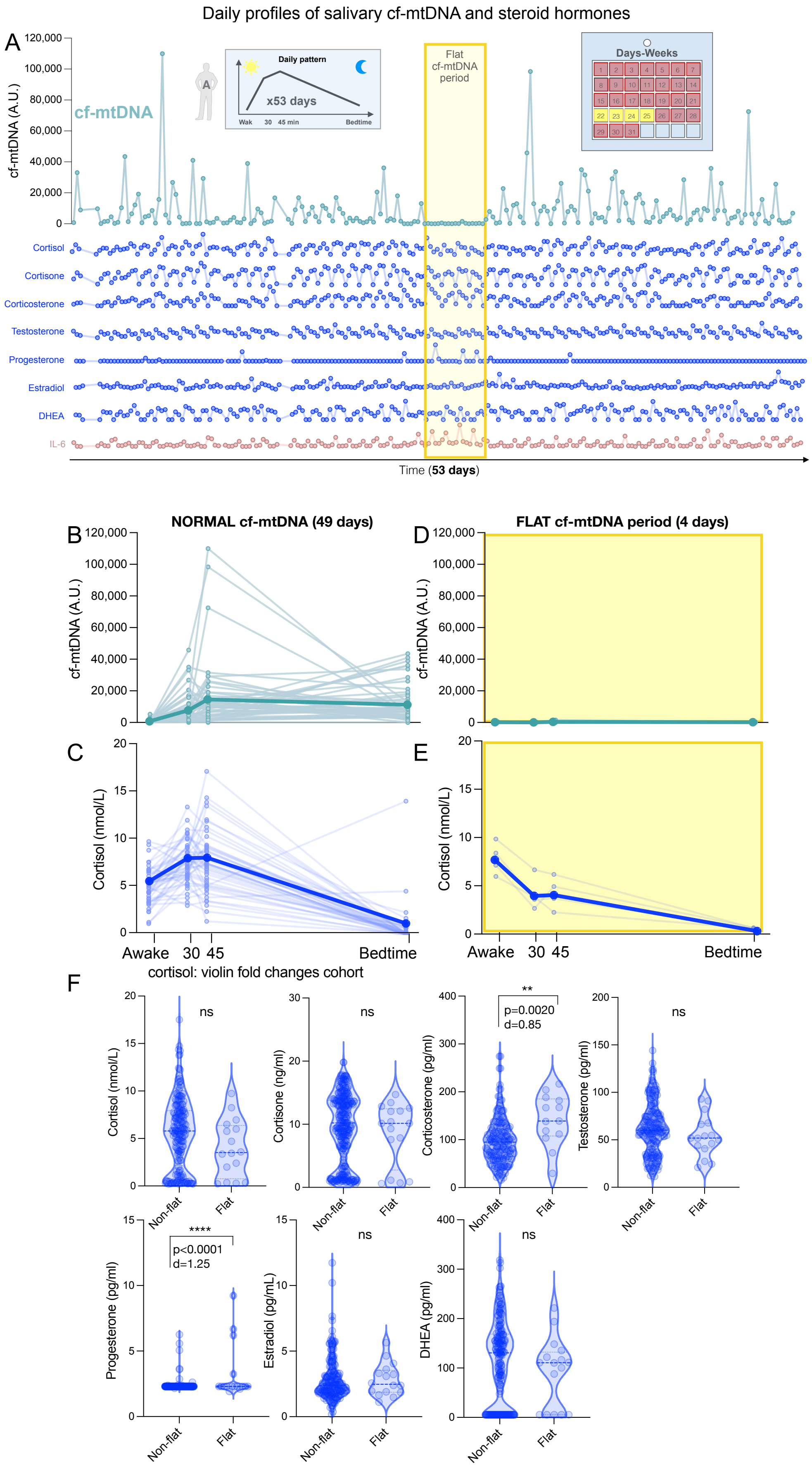


**Figure S8. Salivary cf-mtDNA diurnal variation and its association with steroid hormones.** (**A**) Day-to-day variation of salivary cf-mtDNA and steroid hormones in participant A. (*Top*) Saliva cf-mtDNA collected at awakening, 30 and 45 min after awakening, and bedtime over 53 consecutive days (n=212 observations). A naturally-occurring 4-day period (when someone in the participant's social circle passed away) showed unusually low cf-mtDNA levels, highlighted in yellow. (*Bottom*) Same time course for steroid hormones. Cortisol, cortisone and corticosterone were elevated on the evening preceding the flat cf-mtDNA period, progesterone was highest during the flat period. Comparison of the diurnal profiles of cf-mtDNA and cortisol for the days showing cf-mtDNA levels variation (n=49) (**B,C**) and for the identified 4-day period where cf-mtDNA levels were abnormally low and flat (**D, E**). Data is shown as means and individual datapoints. (**F**) Comparison of steroid hormones for the days showing (non-flat, n=49) and not showing (flat, n=4) cf-mtDNA levels variation. P-value from unpaired t-test, $p<0.05$*, $p<0.01$**, $p<0.001$***, $p<0.0001$***.

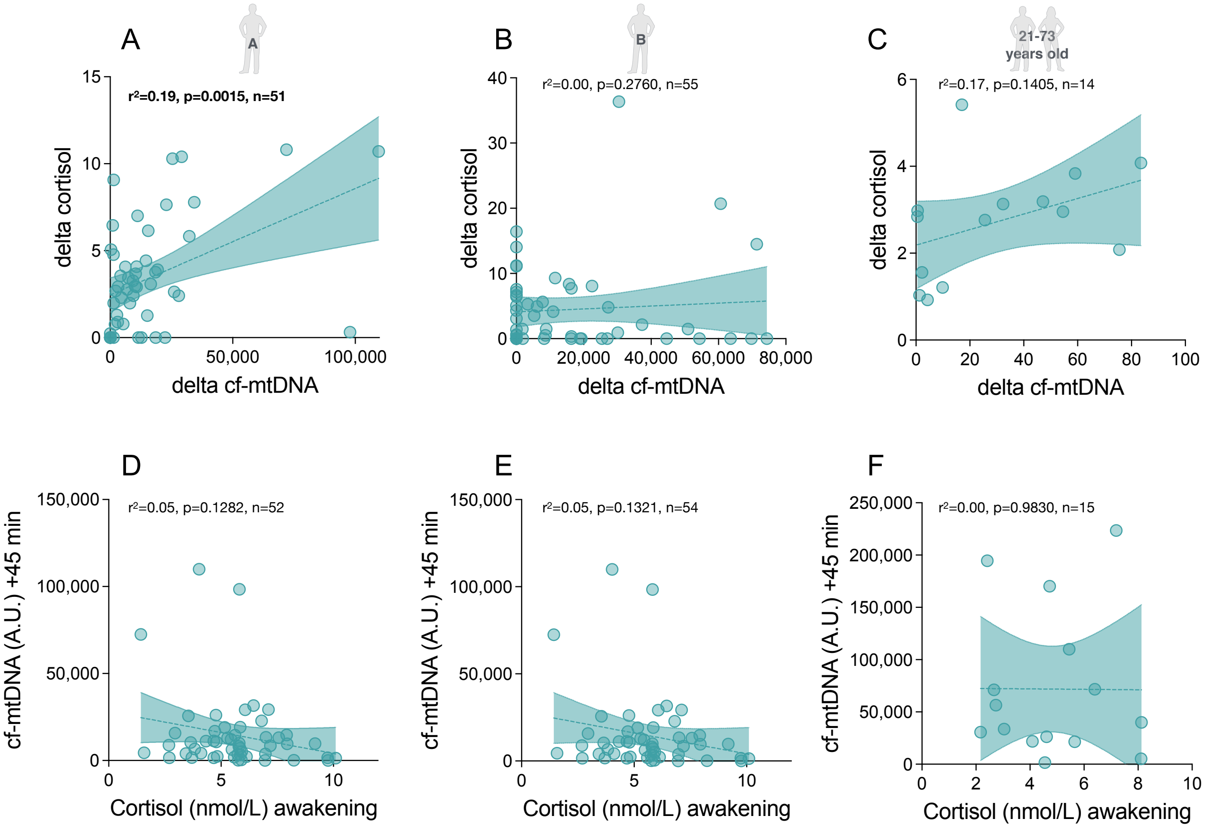


**Figure S9. Salivary cortisol and cf-mtDNA awakening response.** Association between awakening response (highest morning values - awakening) in cortisol and cf-mtDNA levels in (**A**) participant A (n=207 observations), (**B**) B (n=220 observations), and in (**C**) the 4 cohort participants (22 and 31 year old women, 25 and 34 year old men) on 3-4 consecutive days. P-value from simple linear regression. Association between cortisol at awakening and cf-mtDNA +45 min in (**D**) participant A (n=207 observations), (**E**) participant B (n=220 observations), and in (**F**) the 4 cohort participants (22 and 31 year old women, 25 and 34 year old men) on 3-4 consecutive days (n=14 days). P-value from simple linear regression.

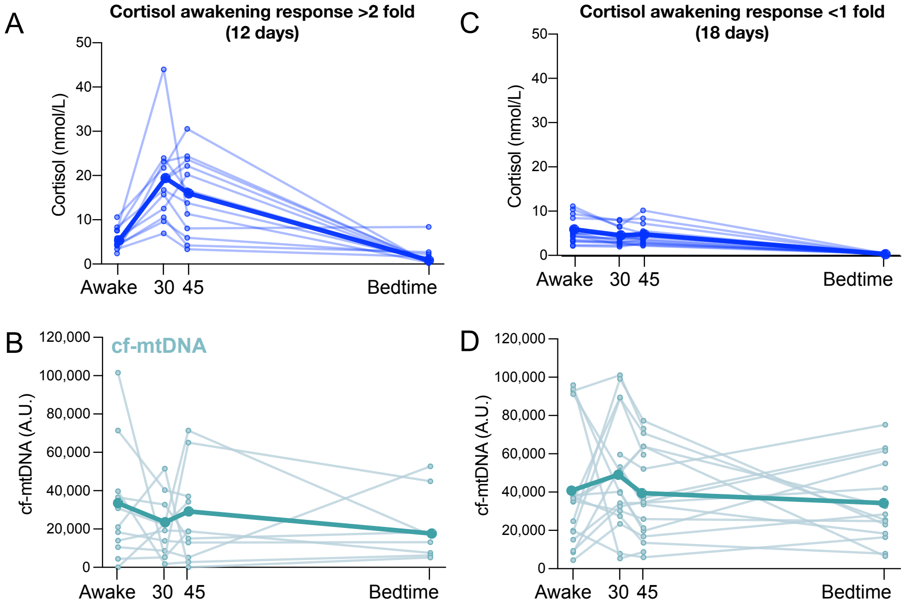


**Figure S10. Salivary cortisol and cf-mtDNA awakening response**. Comparison of the diurnal profiles of (**A**) cortisol and (**B**) cf-mtDNA for the days showing cortisol awakening response from 0 to 30 minutes (fold change >2) (n=12) in participant B. (**C-D**) Same as in A-B for the days for which no cortisol awakening response was observed from 0 to 30 minutes (n=18). Data is shown as means and individual datapoints.

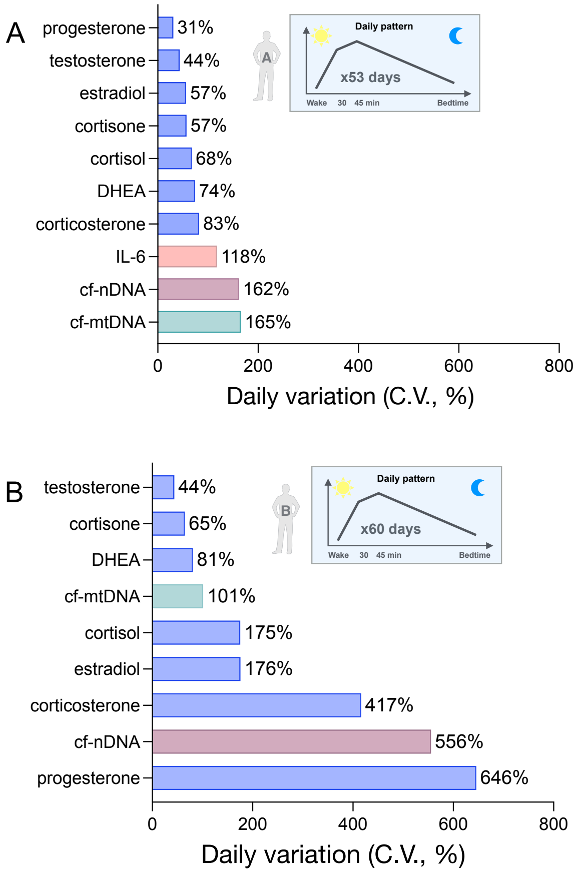


**Figure S11. Salivary cf-DNA and steroid hormones variation.** Coefficient of variation of salivary cf-mtDNA, steroid hormones and IL-6 collected at awakening, 30 and 45 min after awakening, and bedtime in (A) participant A (N=212 timepoints) and (B) participant B (N=220 timepoints). IL-6 levels were not measured in participant B.

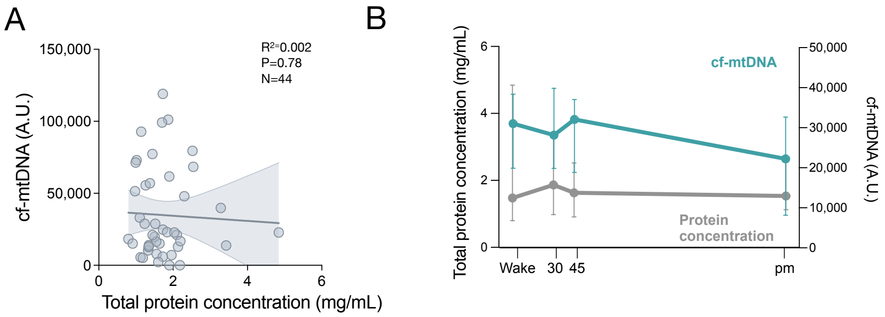


**Figure S12. Saliva protein concentration and cf-mtDNA.** (**A**) Correlation between salivary cf-mtDNA and total salivary protein concentration (N=44 timepoints, Participant B). (**B**) Day-to-day variation of salivary total protein concentration (median (error)) and cf-mtDNA. Data is shown as medians ± SEM.

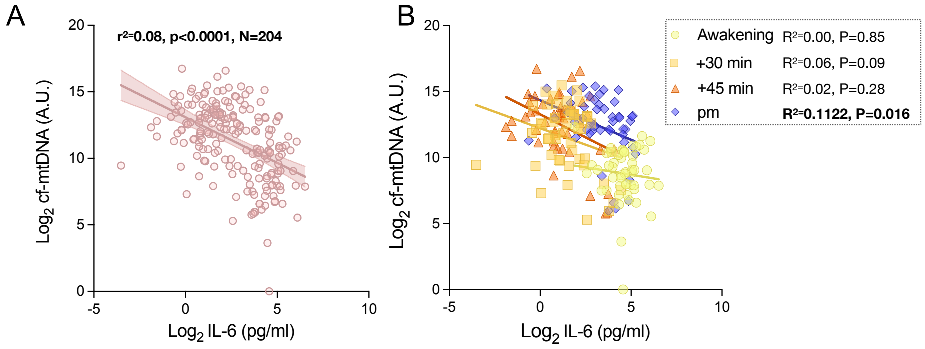


**Figure S13. Association between salivary IL-6 and cf-mtDNA levels.** (**A**) Correlation between salivary IL-6 and cf-mtDNA levels in participant A (n=204 observations). (**B**) Same as in (A), with datapoints color-coded by time of collection.
